# Experimental demonstration that screening can enable the environmental recruitment of a defensive microbiome

**DOI:** 10.1101/375634

**Authors:** Tabitha Innocent, Neil Holmes, Mahmoud Al Bassam, Morten Schiøtt, István Scheuring, Barrie Wilkinson, Matthew I Hutchings, Jacobus J Boomsma, Douglas W Yu

## Abstract

Many animals and plants recruit beneficial microbes from the environment, enhancing their defence against pathogens. However, we have only a limited understanding of the assembly mechanisms involved. A game-theoretical concept from economics, *screening,* potentially explains how a host selectively recruits mutualistic microbes from the environment by fomenting and biasing competition among potential symbionts in such a way that the more likely winners are antibiotic producers. The cuticular microbiomes of *Acromyrmex* leaf-cutting ants inspired one of the first applications of screening theory, and here we simulate this system *in vitro* to test screening. On agar infused with antibacterial metabolites from *Acromyrmex*’s vertically transmitted *Pseudonocardia* bacteria, we show that antibiotic-producing *Streptomyces* bacteria exhibit higher growth rates than do non-antibiotic-producer strains and are more likely to win in direct competition. Our results demonstrate how game-theoretical concepts can provide powerful insight into host-microbiome coevolution.

## Introduction

Many, perhaps most, animal and plant species recruit multiple strains of beneficial microbes from the environment, with one of the most common benefits being defence against pathogens (Kaltenpoth 2009; Barke *et al.* 2011; Seipke *et al.* 2012; Antwis *et al.* 2015; Loudon *et al.* 2016; Duarte *et al.* 2018; Engl *et al.* 2018). It is widely expected that diverse microbiomes provide a more reliable defence against pathogens than do single-species microbiomes, by allowing the application of multi-drug therapy (reviews in Scheuring & Yu 2012; Seipke *et al.* 2012; Antwis & Harrison 2018; Duarte *et al.* 2018; Engl *et al.* 2018). However, despite an abundance of studies on microbiomes, two recent reviews still conclude that “our ability to make predictions about these dynamic, highly complex communities is limited” (Hoye & Fenton 2018), and that “integration between theory and experiments is a crucial ‘missing link’ in current microbial ecology” (Widder *et al.* 2016).

### The Problem of Hidden Information

We have previously shown that game theory from economics provides a powerful shortcut to modelling the coevolution of hosts and their symbionts (Edwards *et al.* 2006; Weyl *et al.* 2010; Archetti *et al*. 2011a). Specifically, a host can be conceptualised as a ‘Principal’ that tries to recruit one or more suitable ‘Agents’ (symbionts) out of all possible Agents (e.g. all microbes in the environment), some or most of which are unsuitable (Archetti *et al.* 2011b; Scheuring & Yu 2012). The challenge is that the Principal does not know which Agents are suitable – that is, *the Agent’s characteristics are hidden*. This is known as the Problem of Hidden Information in economics. The equivalent statement in biology is that in horizontally transmitted symbioses, a host is under selection to recruit mutualistic symbionts out of a species pool that also contains commensals and parasites, but the host is faced with the Partner Choice problem (Bull & Rice 1991) of not being able to recognise which species are mutualistic.

Game theory provides two solutions to the Problem of Hidden Information: *signalling* and *screening* (Archetti *et al.* 2011a; b). Signalling uses the display of costly phenotypes to honestly reveal the quality of potential partners (Spence 1973; Grafen 1990), but it is difficult to envision a cost that can signal cooperativeness *per se* (although see Archetti *et al*. 2016). Also, even if the host is able to discern symbiont qualities, the host might be unable to actively choose amongst the symbionts. Both problems apply to the recruitment of defensive symbionts where the host cannot use *signalling* because it cannot differentiate symbiont lineages before establishment, or has no mechanism for choosing amongst lineages, or both.

### Screening

Here, we present a set of *in vitro* experiments to demonstrate the potential for ant hosts to instead use *screening* to selectively take up antibiotic-producing microbes from the pool of bacteria in the environment. First, we recap how screening works.

Imagine a host faced with multiple potential symbionts differing in their benefit to the host. For simplicity, we assume two types, mutualistic and parasitic, where the latter includes commensals because they can impose opportunity costs on hosts (Yu 2001). It is possible for the host to selectively ‘screen-in’ mutualistic symbionts, provided that the host evolves a ‘demanding environment’, which imposes a cost on colonising symbionts that is easier to endure if they are mutualists. The benefit of enduring the cost is a host-provided reward that is high enough for the mutualist to enjoy a net benefit. (See S5 for a human example of screening.)

Multiple screening mechanisms appear to exist in nature (Archetti *et al.* 2011a; b; see also Ranger *et al.* 2018), and in particular, we have proposed an elaboration of screening in which hosts evolve to foment and bias competition among potential symbionts in such a way that mutualists enjoy a higher probability of being the winners, which we call *competition-based screening* (Archetti *et al.* 2011b). A good example is given by Heil (2013), who studied *Acacia* ant-plants, which can be colonised by ant species that either protect their hostplants (mutualists) or not (parasites). *Acacia* species provide abundant food and housing to incipient ant colonies to promote colony growth and worker activity, and the colony that wins the hostplant is the one whose workers kill off the other incipient colonies. Having numerous, aggressive workers is also the defining characteristic of a mutualist, since such workers attack herbivores. Thus, *the demanding environment is interference competition amongst the ants themselves*, fueled by the hostplant, and the ants best able to endure this cost are the aggressive, ergo mutualistic, species. The benefit of enduring the demanding environment is the high levels of plant-provided reward. This fitness gain, minus the cost of the risk of colony death due to fighting, is presumably greater than the fitness gained by living elsewhere. What makes this an especially satisfying example is that there also exist *Acacia* species that provide low amounts of food and housing, which are regularly colonised by timid, non-defending ants, whereas high-reward *Acacia* species are usually colonised by aggressive ants.

### Attine defensive microbiomes and the Red Queen

We have argued that competition-based screening is consistent with several biological details of the defensive cuticular microbiomes of attine fungus-growing ants (Scheuring & Yu 2012). These microbiomes produce antibiotics that can control specialised crop diseases, in particular mycopathogens in the genus *Escovopsis* (Currie *et al.* 2006), and may also provide brood hygienic protection (Mattoso *et al.* 2012). By biomass, the dominant actinobacterial constituent of attine cuticular microbiomes is the genus *Pseudonocardia*, and there is considerable evidence that attine ants have maintained a long-term, coevolved, and mutualistic relationship with *Pseudonocardia*, via vertical transmission (Cafaro *et al.* 2011; Andersen *et al.* 2013; Steffan *et al*. 2015; Li *et al*. 2018). However, it is recognised that an exclusive relationship between an ant lineage and its vertically transmitted bacterial lineage raises the Red Queen problem (Currie et al. 2006): how can *Pseudonocardia* maintain effectiveness against *Escovopsis weberi*, which reproduces sexually and has evolved countermeasures in the form of two metabolites that are upregulated after infecting the attine fungal cultivar: *melinacidin* and *shearinine*. Both these virulence factors are potent growth suppressors of the *Pseudonocardia* strains (a less restrictive term used in preference to ‘species’ in microbiology) that are hosted by *Acromyrmex* ants, and shearinine also paralyses and kills *Acromyrmex echinatior* workers (Heine et al. 2018).

### Multi-species defensive microbiomes

Multi-species defensive microbiomes are an obvious candidate solution to the Red Queen problem, allowing ants to apply multidrug therapy against *Escovopsis*. Multiple amplicon and culture studies (Kost et al. 2007; Haeder et al. 2009; Sen et al. 2009; Barke et al. 2010; Schoenian et al. 2011, Andersen et al. 2013) have documented the presence of actinomycete genera such as *Streptomyces* and *Amycolatopsis*, which are well-known antibiotic producers (Kaltenpoth 2009; Barka *et al.* 2016; Worsley *et al.* 2018), including an amplicon study reporting *Streptomyces* read frequencies of >5% on eleven field-collected *Acromyrmex echinatior* workers and *Amycolatopsis* read frequencies of 35% and 99% on two *Trachymyrmex zeteki* workers (Andersen et al. 2013). These strains are almost certainly acquired horizontally (i.e. from the environment), but presence alone does not demonstrate that these strains are live and contributing antibiotics. However, Schoenian *et al*. (2011) used liquid-chromatography-mass-spectrometry (LC-MS) to demonstrate that *Streptomyces* cultured from the cuticles of *Acromyrmex* workers (Haeder et al. 2009) synthesise antifungal and antibacterial valinomycins, antimycins, actinomycins, and candicidins, which act synergistically *in vitro* to inhibit *Escovopsis*, thus demonstrating the possibility of multi-drug therapy. Crucially, Schoenian *et al*. also used MALDI imaging to visualise up to several nanograms of valinomycin on the cuticles of *Acromyrmex echinatior* workers, thus demonstrating *in vivo* metabolic activity of *Streptomyces*. Also, Worsley *et al*. (S. Worsley *et al*., *unpubl. data*) have recently used stable isotope probing followed by RNA-sequencing (SIP-RNA) of cuticular bacteria isolated from *Acromyrmex echinatior* workers that had been fed ^13^C-glucose solution to document the uptake of ^13^C by twelve bacterial genera, including *Pseudonocardia* and *Streptomyces*, representing ~40% and ~20% of sequence reads, respectively. In sum, the weight of evidence is shifting toward the conclusion that (at least) *Acromyrmex* recruits and supports a multi-species defensive microbiome, as has been reported for other taxa (see *Introduction*). This now raises the question of why *Acromyrmex* continues to host *Pseudonocardia*.

### Screening applied to the attine microbiome

Newly hatched *Acromyrmex* workers are inoculated with *Pseudonocardia* by their sisters (Poulsen *et al.* 2003b; Andersen *et al.* 2013; Marsh *et al.* 2014), and the monoculture blooms over the workers, then retreats to the propleural plates of the ventral surface after ~25 days, when the workers start foraging (Poulsen *et al.* 2003a; Andersen *et al.* 2013). This is when other actinomycete bacteria can be detected on the ant.

Scheuring and Yu’s (2012) model argues that because *Pseudonocardia* coats the worker cuticle with antibacterials, the cuticle is rendered a demanding environment that we expect antibiotic-producing strains to be better able to endure. This is because antibiotic-producing bacteria must also have antibiotic resistance, or they would commit suicide when producing antibacterials. Thus, the quality that allows potential symbionts to endure this demanding environment (resistance to antibacterials) is correlated with the quality that makes them mutualistic (antibiotic production). This demanding environment is paired with high rewards from the ant’s exocrine glands (Currie *et al.* 2006; Li et al. 2018; S. Worsley *et al*. unpubl data), and the combination is predicted to foment and bias competition in favour of strains that can produce the most effective antibiotics (Scheuring & Yu 2012).

Screening is potentially consistent with an interesting complication discovered by Andersen *et al*. (2013), who showed that *Acromyrmex* colonies hosting one of the native *Pseudonocardia* strains (*P. octospinosus*, also known as ‘Ps1’) are colonised by a more diverse secondary cuticular microbiome than are *Acromyrmex* colonies hosting the other strain (*P. echinatior*, ‘Ps2’) (strains described in Holmes *et al.* 2016). For simplicity, we refer to these as Ps1 and Ps2. Genome analysis has confirmed that Ps1 and Ps2 code for different sets of secondary metabolite clusters (Holmes *et al.* 2016). Screening theory predicts that Ps2 imposes the more demanding environment, by making stronger or more antibacterials, and thus reducing the likelihood of successful secondary colonisation.

### Tests of screening

Experimental tests of mechanisms that could promote selective recruitment are missing from the microbiome literature (Widder *et al.* 2016; Hoye & Fenton 2018). We address this gap with an experiment that tests screening *in vitro.* Using *Pseudonocardia* cultures, we created two types of environment on agar media, emulating the cuticle of *Acromyrmex* ants: *demanding* (where the agar was infused with metabolites from *Pseudonocardia*), and *undemanding* (where *Pseudonocardia* metabolites were absent). We then introduced a variety of bacterial strains representing two classes of colonisers – antibiotic-producing *Streptomyces* (*producers*) and bacteria that do not produce, or produce fewer, antibiotics (for brevity, we call these *non*-*producers*). We first tested the prediction that *producers* are better able to endure the demanding environment (by comparing colony growth rates) than are *non-producers*, and then we directly competed *producers* and *non-producers*.

We show that *producers* grow relatively more quickly on the *demanding* (*Pseudonocardia*-infused) media despite growing relatively more slowly on the *undemanding* (*Pseudonocardia*-absent) media. We also show that *producers* are the superior competitors on the *demanding* media, while *non-producers* are the superior competitors on the *undemanding* media, as predicted by screening theory. Finally, we confirm the assumption that the *producer* strains are relatively more resistant to antibiotics and the prediction that *Pseudonocardia* Ps2 more strongly suppresses bacterial growth. We emphasise that *producer* and *non-producer* are convenience terms, because the *non-producer* strains that we use can produce antibacterials and exhibit resistance, just to a lesser extent than do the *producers*.

## Materials & Methods

We obtained:

(**1**) a set of *Pseudonocardia* Ps1 and Ps2 strains to grow on agar plates. *Pseudonocardia* releases antibacterials and antifungals when grown in culture (Barke *et al.* 2010; Holmes *et al.* 2016; Van Arnam *et al.* 2016);

(**2**) a set of ‘environmental’ *antibiotic-producer* bacterial strains, defined as not having an evolutionary history of growing on the cuticle of *Acromyrmex* ants;

(**3**) a set of ‘environmental’ *non-producer* bacterial strains, defined as not having an evolutionary history of growing on the cuticle of *Acromyrmex* ants; and

(**4)** a set of bacterial strains that were isolated in the cuticular microbiomes of *Acromyrmex* workers and based on their taxonomies, should be classed as *non-producers*. We asked why these bacteria do not competitively exclude the resident antibiotic-*producers*, since they are presumably resistant to *Pseudonocardia* metabolites but do not pay the cost of producing antibiotics. We use the term ‘resident’ only to indicate that we isolated these strains from *Acromyrmex*; we do not imply that these strains are vertically transmitted like *Pseudonocardia*.

For growth media, we used Soya Flour + Mannitol (SFM) agar (20 g soya flour, 20 g mannitol, 20 g agar in 1 L tap water) and lysogeny broth-Lennox (LB-Lennox) media (10 g tryptone, 5 g yeast extract, 5 g NaCl in 1 L deionised water).

### Collection and isolation of bacterial strains

(**1**) Nineteen Ps1 and Ps2 strains were isolated from the cuticles of individual workers of the leaf-cutting ant *Acromyrmex echinatior* (Hymenoptera, Formicidae, Attini). The 19 colonies of these workers were collected in Soberania National Park, Panama, between 2001 and 2014, and reared in climate-controlled rooms at the University of Copenhagen at ca. 25 ºC and 70% relative humidity. Worker propleural plates were scraped with a sterile needle under a stereomicroscope and streak-inoculated across LB-Lennox media under sterile conditions. The plates were then incubated at 30 °C (standard culturing temperature) until visible growth of distinctive white ‘cotton ball’ colonies of *Pseudonocardia* were apparent. These individual colonies were removed under sterile (LAF bench) conditions and inoculated onto fresh plates, a process that was repeated until clean *Pseudonocardia* cultures were isolated. The *Pseudonocardia* isolates were then grown on Mannitol Soya Flour (SFM) media (optimal for actinobacterial growth and spore production), from which spore stock solutions were prepared in 20% glycerol and kept at −20 °C until use. Each isolate was genotyped to Ps1 or Ps2 (Supplementary Information S1). Five strains from each of these two species have previously been genome-sequenced and formally described as different *Pseudonocardia* species by Holmes *et al*. (2016): *P. octospinosus* (Ps1) and *P. echinatior* (Ps2).

(**2**) The 10 environmental *producer* strains (all *Streptomyces*) were taken from general collections in the Hutchings lab (Tables 1, S2).

**Table 1.**
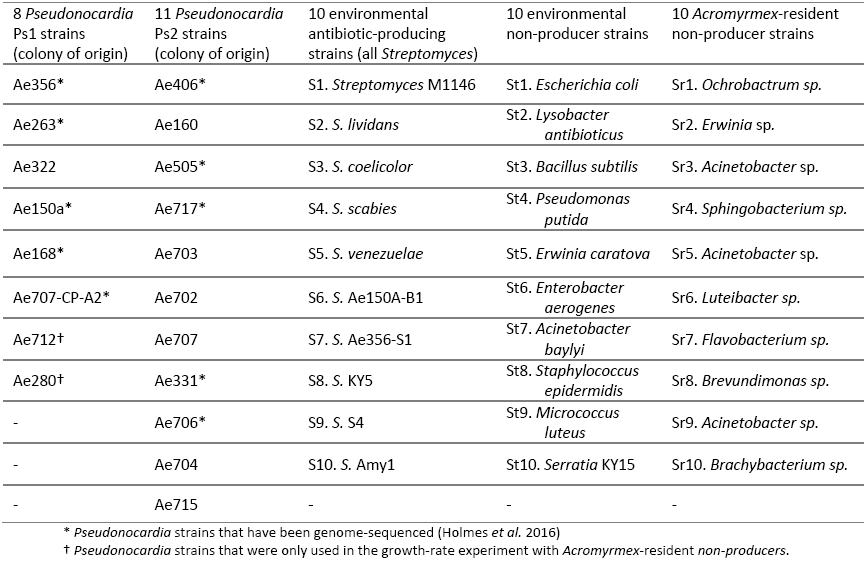
The bacterial strains used in our experiments: the 19 *Pseudonocardia* strains isolated from *Acromyrmex echinatior* (columns 1 and 2); the 10 environmental *antibiotic-producer* strains (column 3); the 10 environmental *antibiotic-non-producer* strains (column 4); and the 10 *Acromyrmex*-*resident, antibiotic‐ non-producer* strains (column 5). Details for all strains not isolated from *Acromyrmex* in Table S2.

(**3**) The 10 environmental *non-producer* strains were obtained from the Hutchings lab (2 strains) and from the ESKAPE suite (8 strains) – a set with varying origins (human skin, soil, etc.) associated with hospital-acquired infections and used to test antibiotic resistance or efficacy in clinical/research settings (Tables 1, S2).

(**4**) The 10 *Acromyrmex*-resident, *non-producer* strains were isolated from *A. echinatior* and *A. octospinosus* ants by culturing from cuticular washes. These strains were 16S-genotyped and where possible assigned to genus by comparison to the NCBI 16S RefSeq database (Supplementary Information S1, Tables 1, S2). Hereafter, for brevity, we refer to the environmental *non-producers* simply as *non-producers* and continue to use the full term ‘resident *non-producers*’ to differentiate them.

### Individual growth-rate experiments

We measured bacterial colony growth rates by growing all *producer* and *non-producer* strains on *Pseudonocardia*-infused and control media.

To create the *Pseudonocardia*-infused media, we grew lawns of the 19 *Pseudonocardia* isolates (Table 1), plating 30 μl on 90 mm SFM agar plates. The control plates were inoculated with glycerol only (20%; identical to that used for making solutions from *Pseudonocardia* strains). We incubated these plates at 30 °C for 6 weeks, which ultimately produced lawns from 17 strains that could be included in the experiments (6 Ps1, 11 Ps2). Once each plate was fully (not necessarily uniformly) covered, we flipped the agar to reveal a surface open for colonisation.

The 10 environmental *producers* and 10 environmental *non-producers* (Table 1) were inoculated onto the plates. Each plate was inoculated with 10 evenly spaced colonies, with 3 replicates, generating 2 invader types x 17 Ps-media-types x 3 replicates = 102 Ps-infused plates and 2 invader types x 3 replicates = 6 control plates, for 1020 treatment and 60 control inoculations. Each strain inoculation used 5 μl of solution (approx. 1 x 10^6^ cells per ml in 20% glycerol), spotted at evenly spaced positions and without coming into direct contact. All plates were incubated at 30 °C for five days, after which photographs were taken.

Images were processed in Fiji software (Schindelin *et al.* 2012; Rueden *et al.* 2017), creating binary negatives (black & white) so automated tools could identify discrete areas of growth (black) and measure growth areas for each invading strain; in the few cases where binary image resolution was insufficient, outlines were added manually before area calculation. 48 *producer*-inoculated and 57 *non-producer*-inoculated treatment measurements were excluded because plate condition had deteriorated to become unscorable or they were contaminated, leaving final sample sizes of 1020-48-57= 915 treatment inoculations and 60 controls.

The second growth-rate experiment compared the 10 *Acromyrmex*-resident, *non-producer* strains with 9 of the environmental *producer* strains (1 of the 10 inoculations failed to grow). All 19 *Pseudonocardia* strains grew sufficiently to be included in this experiment, and each plate was again inoculated with 10 or 9 evenly spaced colonies. Starting sample sizes were therefore 2 invader types x 19 Ps-media-types x 3 replicates = 114 Ps-infused plates and 2 x 3 = 6 control plates, for 1083 treatment and 57 control inoculations. 50 *producer* and 20 *non-producer* treatment measurements were excluded for the same reasons as above, leaving final sample sizes of 1083-50-20=1013 treatment and 57 control inoculations, scored as above.

### Pairwise competition experiment

We then tested whether *producers* could win in direct competition against *non-producers*. We co-inoculated pairs of environmental *producers* and *non-producers* on *Pseudonocardia*-infused media and on control media (prepared as in the individual growth-rate experiment), and we measured the outcome of competition as a win, loss, or draw. The screening model (Scheuring & Yu 2012) predicts that the *producer* strains is more likely to win when growing on *Pseudonocardia*-infused media.

To keep the number of tests manageable, we used two combinations of *Pseudonocardia*-infused media and *Streptomyces*: *Pseudonocardia* Ps1 (strain Ae707) **+** *Streptomyces* S8 and *Pseudonocardia* Ps2 (strain Ae717) **+** *Streptomyces* S2 (Table 1). These combinations were representative of the results from the growth-rate experiment: the two *Streptomyces* strains grew more slowly than most of the *non-producers* on control media and either near the median growth rate of (S8) or faster than (S2) the *non-producers* on the media infused by these *Pseudonocardia* strains.

We competed the two *Streptomyces* strains (S2, S8) against the 10 environmental *non-producer* strains (20 pairings), using *Streptomyces* spore titrations consisting of 10^6^ spores per ml for each strain in 20% glycerol. The *non-producer* strains were grown overnight in 10 ml LB-Lennox, subcultured with a 1:100 dilution into 10 ml of fresh LB-Lennox, and grown at 37 °C for 3-4 hours, after which OD_600_ was measured, assuming that OD_600_ = 1 represented 8 × 10^9^ cells. Similar dilutions of 10^6^ cells per ml were made for each *non-producer* strain in 20% glycerol, after which *producer* and *non-producer* dilutions were mixed 1:1. Each pair of *producer* and *non-producer* was co-inoculated as a mixture of 20 µl (10^4^ spore-cells of each) on the designated *Pseudonocardia*-infused media (set up as above) with 5 replicates per pairing. We used 150 plates for the S8 experiment (including 100 control plates; 10 replicates per pairing) and 100 plates for the S2 experiment (including 50 control plates; 5 replicates per pairing). Plates were incubated at 30 °C for 5 days and then photographed, after which images were scored with respect to the *producer* as: win (dominant growth), draw (both strains growing with no clear dominant), or loss (little or no visible growth), with reference to images of each strain grown alone on control media to minimise observer bias. One plate’s outcome was too ambiguous to score and was omitted. All plates were independently scored by two observers, one using photos, which produced datasets giving the same statistical results. We report the direct observer’s scores.

### Antibiotic-resistance profiles

Our key assumption is that *producer* strains (here, all *Streptomyces*) are relatively better at resisting antibiotics, because this correlation is what confers competitive superiority on a toxic host surface. We tested this assumption by growing the 10 environmental *producer* strains and the 10 environmental *non-producer* strains (Table 1, S2) in the presence of 8 different antibiotics, representing a range of chemical classes and modes of action. Antibiotics were added to 1 ml of LB-Lennox medium in a 24-well microtitre plate at 6 different concentrations. The relative concentration range was the same for each antibiotic, although actual concentrations reflected activity (Table S3). *Producers* and *non-producers* were inoculated onto the plates and incubated at 30 °C for 7 days, then photographed (Table S3). Lowest Effective Concentration (LEC, lowest concentration with inhibitory effect) and Minimum Inhibitory Concentration (MIC, lowest concentration with no growth) scores were assigned on a Likert scale of 1–6, where 1 was no resistance and 6 was resistance above the concentrations tested (adapted from generalised MIC methods; reviewed by Balouiri *et al.* 2016). The Likert scale is ordinal, so we analysed using a non-parametric test.

### Justification of experimental-design choices

Petri-dish experiments are analogous to greenhouse experiments in plant ecology in that they trade off realism for improved inference of cause and effect, but microbial experiments face the additional challenge of having to use macroscopic elements to mimic microscopic interactions. For instance, we stage competition between pairs of strains at the macroscopic scale because it is likely to be the more realistic representation of competition at the microscopic scale, since interactions are local and colonising spores from the air arrive slowly and settle randomly. These considerations cause the instantaneous probability of interaction between strains to be low, and the losing strain to be excluded before a new invader arrives in the same position, rendering competition effectively pairwise in real microbiomes. Pairwise competition is also more conservative in that it should maximise the competitiveness of each strain competed against *Streptomyces*. If multiple *non-producer* strains were mixed and inoculated together, they would also compete against each other (i.e. ‘an all-out melee’), diminishing the competitiveness of any given strain against *Streptomyces*.

We also conservatively chose not to replenish nutrients after growing *Pseudonocardia* to make the demanding plates, because increased food resources favour *producers*, since antibiotic synthesis and resistance are costly (Scheuring & Yu 2012). Finally, we included ESKAPE strains as *non-producers* because they grow well in lab conditions, thus removing an artefactual cause of competitive inferiority.

### Statistical analyses

Initial models revealed correlated residuals by *Pseudonocardia* strain and inoculant strain, so we used mixed-effects models to incorporate strain identities as random intercepts, and we nested plate replicates within *Pseudonocardia* strain. Statistical significance was determined by term deletion, and final-model residuals were approximately homoscedastic and uncorrelated within random-factor levels. Analyses were carried out in *R* 3.4.4 (R Core Team 2017) with *lme4* 1.1-17 (Bates *et al.* 2015), *tidyverse* 1.2.1 (Wickham 2017), and *RColorBrewer* 1.1-2 (www.colorbrewer.org, accessed 4 Mar 2018). The *R* scripts and data are at github.com/dougwyu/screening_Innocent_et_al and in S4.

## Results

### Individual growth-rate experiments

As predicted by the screening model, the environmental *non-producers* grew more quickly on the undemanding control media while *producers* grew more quickly on the demanding *Pseudonocardia*-infused media, producing a highly significant statistical interaction effect (Fig. 1A). There was also a significant main effect of *Pseudonocardia* genotype, with both *non-producers* and *producers* exhibiting a lower growth rate on Ps2-infused media.

**Figure 1.**
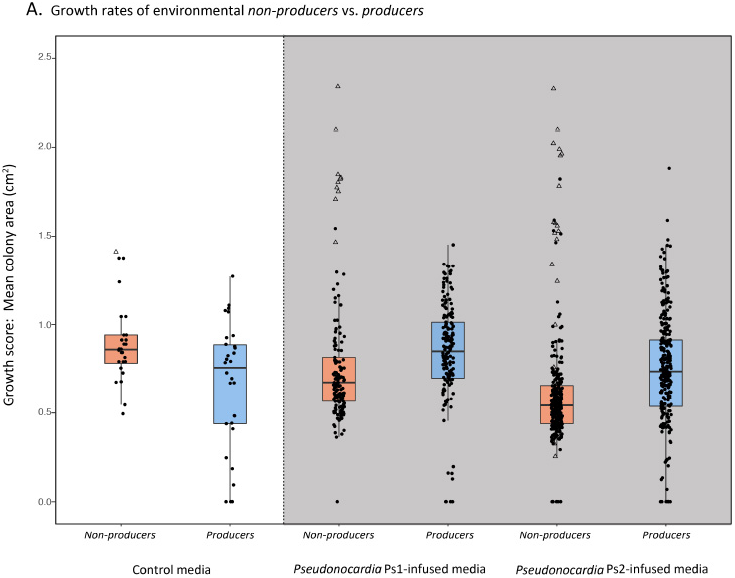
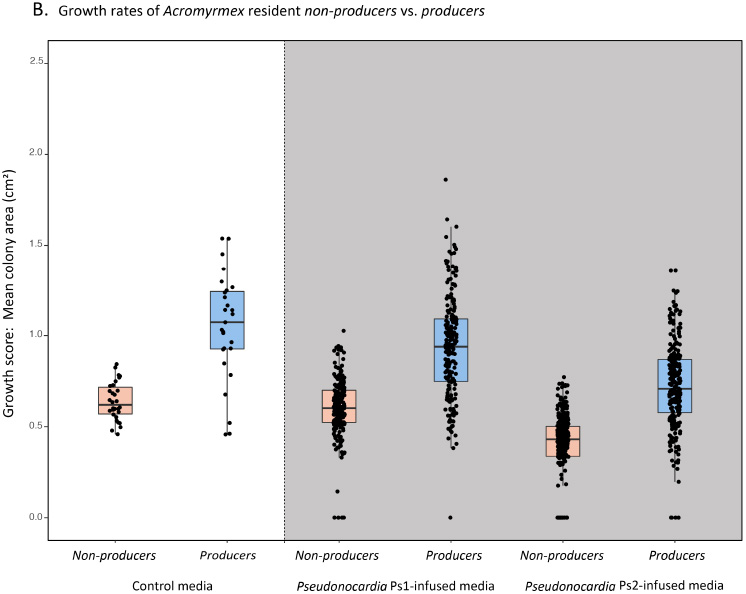
Individual growth-rate experiments. Bacterial colony sizes after 5 days at 30 ºC, with the boxplots indicating medians ± one quartile. The white section shows growth rates on control media, and the grey section shows growth rates on the *Pseudonocardia*-infused media. Coral boxes represent *non-producer* strains, and blue boxes represent *producer* strains. For analysis, a linear mixed-effects model, including *Pseudonocardia* strain (`Ps.strain`, 17 groups) and inoculated bacterial species (`Inv.strain`, 20 groups) as random intercepts, was used to test for interaction and main effects of growth media (`Control vs. Ps1.infused vs. Ps2.infused`) and antibiotic production (`Non.producers vs. Streptomyces`): (*lme4::lmer*(Growth.score ~ Media * Antibiotic.production + (1|Ps.strain/Plate) + (1|Inv.strain)). **A.** Environmental strains. There was a highly significant interaction effect, such that environmental non-producer strains grow more rapidly on control media, while *Streptomyces* grow more rapidly on *Pseudonocardia*-infused media (n = 975, χ^2^ = 45.86, df = 2, p < 0.0001). There was also a highly significant main effect of *Pseudonocardia* genotype, with growth being lower on Ps2-infused media (n = 915, ^2^ = 24.55, df = 1, p < 0.0001, control-media data omitted for this analysis). One non-producer strain (*Staphylococcus epidermidis*) grew more rapidly than all other strains (open triangle points), demonstrating the need to control for correlated residuals by strain. The y-axis has been truncated at 2.5 for clarity (the full figure is in Supplementary Information). **B.** *Acromyrmex*-resident, *non-producer* strains. There was no interaction effect (χ^2^ = 2.64, df = 2, p = 0.27), but both main effects were highly significant. The *non-producers* grow more slowly on all media types (χ^2^ = 20.96, df = 1, p < 0.0001), and bacterial growth was generally slower on Ps2-infused media (χ^2^ = 21.43, df = 1, p < 0.0001, control-media data omitted).

The resident *non-producers* isolated from cuticular microbiomes had significantly slower growth rates overall, even on the non-toxic control media (Fig. 1B). This explains why they are unable to outcompete *producer* strains but raises the question of why these *non-producer* species persist at all. We return to this in the Discussion.

### Pairwise competition experiment

When grown on the demanding *Pseudonocardia*infused media, the *producer* strains were much more likely to win in direct competition against the *non-producer* strains (Fig. 2). Draws were more common on the Ps1-infused plates.

**Figure 2.**
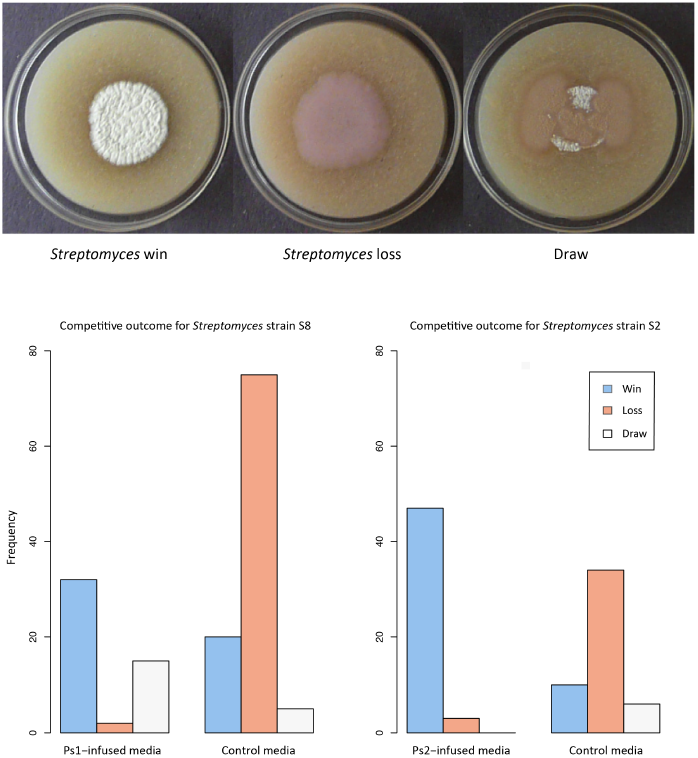
Pairwise competition experiment, scoring whether or not *producers* prevail. **A**. Representative images of agar plates at five days post-inoculation showing examples of the three competitive outcomes: Win (S2 vs. St3 on Ps2 media), Draw (S2 vs. St3 on control media), and Loss (S8 vs. St3 on control media; strain details in Table 1). **B**. Barcharts of competitive outcomes for the two *Streptomyces producer* strains (S8 and S2; Table 1), each pairwise co-inoculated with ten non-antibiotic-producer strains on *Pseudonocardia*-infused media and on control media. For analysis, draw outcomes were omitted, and a general linear mixed-effects model, including non-antibiotic-producer strain as a random intercept (10 groups), was used to test for an effect of the medium term (Control vs. Ps1/2-infused) on competitive outcome (Win vs. Loss) ((*lme4::glmer*(outcome ~ medium + (1|non.producer.strain), family = binomial)). Significance was estimated using term deletion. In both experiments, *producer* strains (*Streptomyces*) were more likely to be competitively superior when grown on the *Pseudonocardia*-infused media (Left: n = 129, χ^2^ = 103.6, df = 1, p < 0.0001; Right: n = 94, χ^2^ = 87.9, df = 1, p < 0.0001).

### Antibiotic-resistance profiles

The screening model assumes that the *producer* strains are more resistant to antibiotics than are the *non-producer* strains. We confirmed this assumption; the Lowest Effective Concentration (LEC) and the Minimum Inhibitory Concentration (MIC) scores were both greater for the *producer* strains (Fig. 3). We also demonstrated that the *resident non-producer* strains had high levels of antibiotic resistance (Fig. 3), as expected, given that they were isolated from ant cuticles (Fig. 1B).

**Figure 3.**
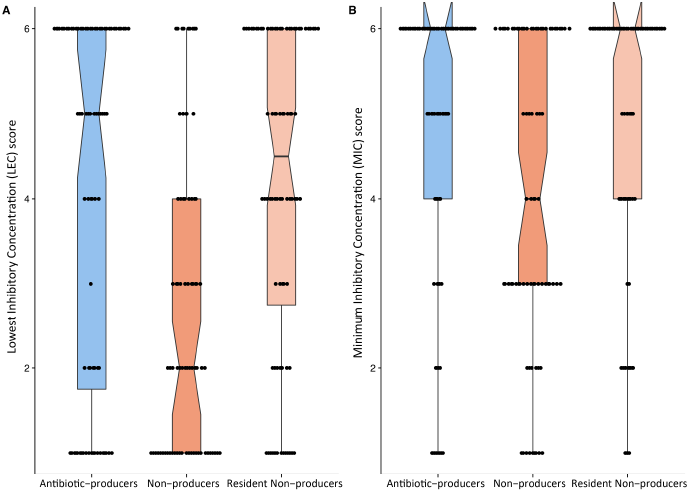
Antibiotic resistance profiles for *producer*, *non-producer*, and *resident non-producer* strains (Table 1). Boxplots indicate medians (notches) ± one quartile. For analysis, we calculated each strain’s mean growth score across the eight tested antibiotics (reducing from n = 155 to n = 20; details in S3). *Producers* showed higher levels of resistance than did *non-producers* for both measurements: Wilcoxon two-sided test `(wilcox.test)`, W = 94.5, p = 0.0017 for LEC (**A**) and W = 80, p = 0.0253 for MIC (**B**), after correction for two tests (*p.adjust*(method=“fdr”)). *Producers* and *Resident Non-producers* showed no difference in resistance levels (p = 0.44 and 0.25). The ‘rabbit ears’ in **B** indicate that the medians are also the highest values.

## Discussion

We used *in vitro* experiments to test key predictions and assumptions of a screening model (Scheuring & Yu 2012) that explains how leaf-cutting ants could use a vertically transmitted symbiont, *Pseudonocardia*, to selectively recruit additional antibiotic-producing bacterial strains out of the large pool of potential bacterial symbionts present in the environment.

We showed that *producer* strains grow relatively faster than do *non-producer* strains on *Pseudonocardia*-infused toxic media (Fig. 1A). Since the *non-producers* have faster growth on the non-toxic control plates, we attribute the *producers’* relatively faster growth on toxic media to greater inhibition of *non-producers* by *Pseudonocardia* metabolites, consistent with the large number of antibacterials documented in *Pseudonocardia*’s genome (Holmes *et al.* 2016). This interpretation is supported by the experiment showing that the *producer* strains are more resistant to a wide range of antibiotics than are the *non-producer* strains (Fig. 3) and is consistent with the result that *Pseudonocardia* Ps2 is a stronger suppressor of bacterial growth (Fig. 1A), since Ps2-hosting *Acromyrmex* workers host lower-diversity (and more Ps2-dominated) cuticular microbiomes (Andersen *et al*. 2013).

Most importantly, the pairwise competition experiment confirmed that growing on *Pseudonocardia*-infused media renders *producer* strains competitively superior (i.e. they usually win) or at least resistant to exclusion (they more often draw than lose, especially on the less-toxic Ps1-infused media) (Fig. 2). This experiment mimics the life-history stage of *Acromyrmex* workers after their full-body *Pseudonocardia* bloom has retreated to the propleural plates (and possibly into the exocrine glands) and they have begun to forage outside the nest, where they are exposed to bacterial colonists (Poulsen *et al.* 2005; Andersen *et al.* 2013; Marsh *et al.* 2014; Andersen *et al.* 2015).

Taken together, these results show how a vertically transmitted, antibiotic-producing symbiont like *Pseudonocardia* can be used by the ant host to create a demanding environment that biases competition in favour of antibiotic-producers, favouring their successful colonisation. In other words, vertical transmission of one mutualist can enable the successful horizontal transmission of other mutualists. Vertical transmission and horizontal transmission are therefore not competing explanations, consistent with the two transmission modes being found together in other symbioses (Ebert 2013; Bourguignon *et al.* 2018; da Costa & Poulsen 2018; Ivens *et al.* 2018). Under other circumstances, mutualistic recruitment via screening may also be achievable without a vertically transmitted symbiont; Duarte *et al*. (2018) have argued that a host might be able to create a suitably demanding environment by evolving the right set of chemical exudates.

We emphasise that the outcomes of the pairwise competition experiment are the *emergent* outcomes of a complex interplay of mechanisms and stochasticity, including the production, export, diffusion, and decay of a diverse set of antibacterials being blocked, decomposed, neutralised, sequestered, and re-exported by a diverse set of resistance mechanisms, spatially (and possibly temporally) heterogeneous resource supply by the host, and community bistability via space taking caused by stochastic orders of arrival, playing out across innumerable local interactions. A mechanistic model incorporating these complications (Boza *et al*. 2018) still results in the same emergent outcomes as in the original phenomenological model (Scheuring & Yu 2012). Based on this modelling, our best (and still simplified) guess of what is going is that the *Streptomyces* strains are relatively more efficient than the *non-producer* strains at resisting toxic *Pseudonocardia* antibacterials (Fig. 3). This efficiency advantage lets *Streptomyces* devote a larger proportion of available resources toward colony growth (Fig. 1A) and toward production of their own antibacterials (e.g. Schoenian *et al*. 2011), the combined effect of which lets *Streptomyces* capture open space more rapidly than its competitors. This scenario does not require the screened-out bacteria to have zero ability to synthesise and resist antibacterials, just less ability.

We note that the ant can employ *Pseudonocardia* as a screening mechanism regardless of whether *Pseudonocardia*’s fitness is aligned with the ant, because the ant controls where and when host resources are provided, thus governing *Pseudonocardia*’s spatial distribution and biomass. We also note that since virgin *Acromyrmex* queens are inoculated upon hatching by nurse workers that maintain *Pseudonocardia* monocultures, the acquisition of other symbionts by foraging workers should not disrupt verticality in the transmission of *Pseudonocardia*.

Acromyrmex-*resident non-producers*. – We also ran a growth-rate experiment with a set of *resident non-producer* strains that had been isolated from *Acromyrmex* ant cuticles (Fig. 1B) and found that they generally had low growth rates, making it unlikely that they can competitively exclude *producers*. It seems surprising that we found these *non-producer* strains at all, but we note that the fact that we could isolate them does not imply that they are abundant. Resident *non-producer* taxa are only found on some (not all) field-sampled *Acromyrmex* workers (Andersen *et al.* 2013; T. Innocent *et al*., *unpubl. data*) and generally below 1% as a proportion of amplicon read number. They may only be transient environmental contaminants, since many were isolated via whole-ant washes, or they might be persisting via non-equilibrium dynamics. In simulation models (Boza et al. 2018), we have found that if bacterial competition is limited to nearest neighbours and antibiotics slowly diffuse across the ant surface, even non-antibiotic-resistant strains can sometimes persist for a long time.

It remains an open question why three different *Acromyrmex* species in Panama all host the same two *Pseudonocardia* strains, Ps1 and Ps2 (Andersen *et al.* 2013). The genomes of Ps1 and Ps2 strains each encode a single distinct nystatin-like antifungal, plus variable and partially segregated sets of secondary metabolite biosynthetic gene clusters with predicted antibacterial functions (Holmes *et al.* 2016). These two *Pseudonocardia* species are almost never found in the same colony even though they are encountered at approximately equal rates in field colonies (Poulsen *et al.* 2005). We hypothesise that the difference between Ps1 and Ps2 in microbiome invasibility (Fig. 1A) reflects balancing selection over a tradeoff between stronger endogenous production of antimicrobials early in worker life, against the recruitment of a more diverse set of antibiotic producers later in life.

Comparable studies to ours using other ant species will be needed to test the generalisability of screening to other attine ant lineages that maintain actinobacterial cuticular microbiomes, including species that appear to have cuticular microbiota with abundant actinomycete genera other than *Pseudonocardia* (Sen *et al.* 2009; Andersen *et al.* 2013; Meirelles *et al.* 2014). Our results suggest that cuticular microbiomes and host resource provisioning can be simulated on agar plates adequately enough for experimentation, providing a tractable method with which to test microbiome assembly mechanisms, eventually leading to *in vivo* experiments.

Such studies might also shed light on why *Atta* and some other genus-level branches of the attine fungus-growing ants have secondarily lost their cuticular microbiomes, a phenomenon that remains poorly understood. The screening requirement to pair a demanding environment with a high reward imposes nontrivial costs on hosts, and in some cases, this cost might have tipped the cost-benefit ratio in favour of chemical pest control over biological control (Fernandez-Marin *et al.* 2009; 2015).

### Toward a general game theory of hosted microbiomes

Hosted microbiomes are coevolutionary metacommunities in which most of the species interactions are difficult to observe. There are numerous calls to use ecological theory to understand microbiomes (Widder *et al.* 2016; Hoye & Fenton 2018), but we propose that game theory may be more promising for producing testable predictions. Screening (Archetti *et al.* 2011a) and password signalling (Archetti *et al.* 2016) are candidate solutions to the Problem of Hidden Information. When hosts evolve to use screening, the emergence of demanding environments can be thought of as an exercise in applied ecology, where the host evolves mechanisms to foment and bias competition by modifying resource and toxicity levels to encourage the assembly of mutualistic microbial communities (see Foster *et al.* 2017; Duarte *et al.* 2018 for similar perspectives).

## Acknowledgments

The work was supported by the European Research Council (ERC Advanced grant 323085 to JJB), a Marie Curie Individual European Fellowship (IEF grant 627949 to TI), NERC grants NE/J01074X/1 to MH and DY and NE/M015033/1 and NE/M014657/1 to MH, DY and BW. D. W. Yu was supported by the National Natural Science Foundation of China (41661144002, 31670536, 31400470, 31500305), the Key Research Program of Frontier Sciences, CAS (QYZDY-SSW-SMC024), the Bureau of International Cooperation project (GJHZ1754), the Ministry of Science and Technology of China (2012FY110800), the University of East Anglia, the State Key Laboratory of Genetic Resources and Evolution at the Kunming Institute of Zoology, and the University of Chinese Academy of Sciences. IS is supported by National Research, Development and Innovation Office of Hungary (GINOP 2.3.2.-15-2016-0057). We thank the Smithsonian Tropical Research Institute for logistical help and facilities for all work in Gamboa, Panama, and the Autoridad Nacional del Ambiente y el Mar for permission to sample and export ants. We thank Ryan Seipke and Gergely Boza for input in developmental stages of the work; Elaine Patrick and Sylvia Matthiesen for lab and logistics support; Michael Poulsen, Saria Otani and Panos Sapountzis for useful technical advice and discussion. The authors declare no conflicts of interest.

## Data accessibility statement

The R scripts and data for analyses are posted at github.com/dougwyu/screening_Innocent_et_al.

## Author declaration

DY, IS, JJB, MH conceived the research, and DY, MH, JJB, TI, NH, MAB, BW and IS designed the experiments. TI, MAB and MS isolated strains. TI and NH carried out the experiments. DY performed the statistical analysis. DY, JJB, TI and NH wrote the manuscript, with comments from all other authors.

## S1. Genotyping of *Pseudonocardia octospinosus* (Ps1) and *P. echinatior* (Ps2) strains, and *Acromyrmex* resident *non-producer* strains

We confirmed the *Pseudonocardia* genotype of the 19 strains – *P. octospinosus* (Ps1) or *P. echinatior* (Ps2) – using material from a parallel study (T. Innocent *et al*., in prep) where DNA extractions were carried out using the Qiagen Blood & Tissue DNA kit (following the manufacturers’ protocol, with an additional step of vortexing each sample for 30 seconds with 0.1 mm glass beads at the first stage; each sample was re-eluted in 100µl AE elution buffer).

We amplified part of the elongation factor EF-Tu gene using the *Pseudonocardia*specific primers 52F and 920R (Ludwig *et al.* 1993; Poulsen *et al.* 2005). We ran 20 µL PCR reactions using the VWR 2x PCR Mix (VWR, USA) and with the following conditions: 96°C for 2.5 minutes; 40 cycles of 94°C for 45s, 56°C for 50s, 72°C for 2 minutes; and a final extension period of 10 minutes at 72°C. Following PCR amplification, confirmed by observing a single band on 2% agarose gel, DNA was purified using a MSB Spin PCRapace kit (STRATEC Molecular GmbH). All samples were prepared for sequencing using a Mix2Seq kit (Eurofins genomics) and sent to and directly sequenced at MWG Eurofins (MWG, Germany).

We aligned all sequences, and compared to existing Ps1 and Ps2 consensus sequences (Ps1, [GenBank: DQ098127]; Ps2, [GenBank: DQ098133]), using MUSCLE implemented in Geneious R7.1, to assign each strain to one or other species.

The 10 *Acromyrmex*-resident, *non-producer* strains were 16S-genotyped, with strains first grown on agar plates, added to 100 µl 50% DMSO and heated 65°C for 20 minutes. We then amplified 16S rDNA from this material using the primers 28F (5′-GAGTTTGATCNTGGCTCAG-3′) and 519R (5′- GWNTTACNGCGGCKGCTG-3′). We ran 50 µL PCR reactions using BIOTAQ^TM^ Red DNA Polymerase Mix (Bioline) and with the following conditions: 95°C for 1 minute; 30 cycles of 95°C for 15s, 55°C for 15 s, 72°C for 15s; and a final extension period of 2 minutes at 72°C.

Following PCR amplification, confirmed by observing a single band on 1% agarose gel, DNA was purified using a QIAquick Gel Extraction kit (QIAGEN). All samples were prepared for sequencing using a Mix2Seq kit (Eurofins genomics) and sent to and directly sequenced at MWG Eurofins (MWG, Germany).

## S2. Source information for all experimental strains

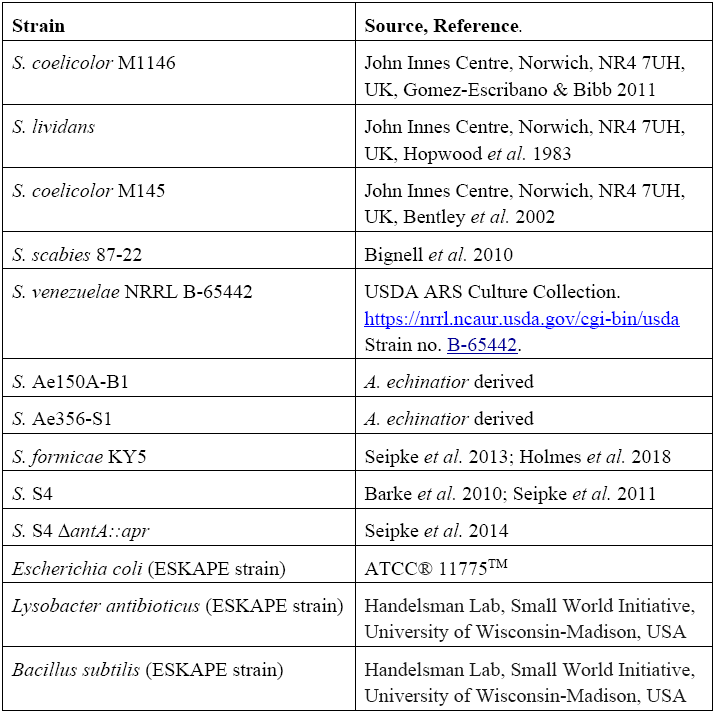
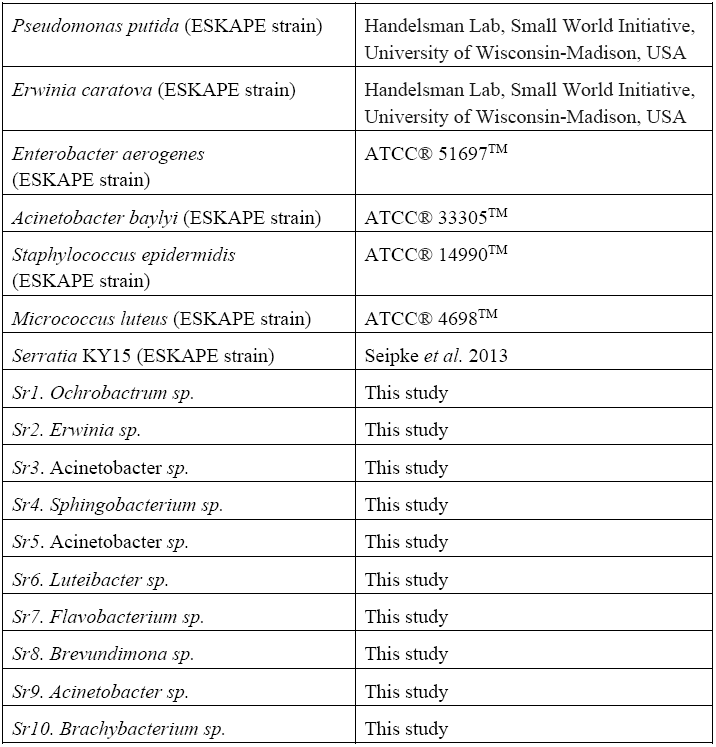

| Strain | Source, Reference. |
| --- | --- |
| *S. coelicolor* M1146 | John Innes Centre, Norwich, NR4 7UH, UK, Gomez-Escribano & Bibb 2011 |
| *S. lividans* | John Innes Centre, Norwich, NR4 7UH, UK, Hopwood *et al.* 1983 |
| *S. coelicolor* M145 | John Innes Centre, Norwich, NR4 7UH, UK, Bentley *et al.* 2002 |
| *S. scabies* 87-22 | Bignell *et al.* 2010 |
| *S. venezuelae* NRRL B-65442 | USDA ARS Culture Collection.<br>https://nrrl.ncaur.usda.gov/cgi-bin/usda<br>Strain no. B-65442. |
| *S.* Ae150A-B1 | *A. echinator* derived |
| *S.* Ae356-S1 | *A. echinator* derived |
| *S. formicae* KY5 | Seipke *et al.* 2013; Holmes *et al.* 2018 |
| *S.* S4 | Barke *et al.* 2010; Seipke *et al.* 2011 |
| *S.* S4 $\Delta antA::apr$ | Seipke *et al.* 2014 |
| *Escherichia coli* (ESKAPE strain) | ATCC® 11775™ |
| *Lysobacter antibioticus* (ESKAPE strain) | Handelsman Lab, Small World Initiative, University of Wisconsin-Madison, USA |
| *Bacillus subtilis* (ESKAPE strain) | Handelsman Lab, Small World Initiative, University of Wisconsin-Madison, USA |
| *Pseudomonas putida* (ESKAPE strain) | Handelsman Lab, Small World Initiative, University of Wisconsin-Madison, USA |
| *Erwinia caratova* (ESKAPE strain) | Handelsman Lab, Small World Initiative, University of Wisconsin-Madison, USA |
| *Enterobacter aerogenes* (ESKAPE strain) | ATCC® 51697™ |
| *Acinetobacter baylyi* (ESKAPE strain) | ATCC® 33305™ |
| *Staphylococcus epidermidis* (ESKAPE strain) | ATCC® 14990™ |
| *Micrococcus luteus* (ESKAPE strain) | ATCC® 4698™ |
| *Serratia* KY15 (ESKAPE strain) | Seipke *et al.* 2013 |
| Sr1. *Ochrobactrum* sp. | This study |
| Sr2. *Erwinia* sp. | This study |
| Sr3. *Acinetobacter* sp. | This study |
| Sr4. *Sphingobacterium* sp. | This study |
| Sr5. *Acinetobacter* sp. | This study |
| Sr6. *Luteibacter* sp. | This study |
| Sr7. *Flavobacterium* sp. | This study |
| Sr8. *Brevundimona* sp. | This study |
| Sr9. *Acinetobacter* sp. | This study |
| Sr10. *Brachybacterium* sp. | This study |

## S3. Detailed methods for antibiotic MIC tests

*Antibiotic-producer strains*: From spore titrations of *Streptomyces*, 1 x 10^6^ spores of each strain were plated onto SFM media with each of the eight antibiotics at six concentrations (Table S3).

*Environmental non-producer and* Acromyrmex*-resident non-producer strains*: These were grown over 1-2 nights in 10 ml Lennox; sub-cultured with a 1:100 dilution into a fresh 10 ml Lennox; grown again at 37 °C for 3-4 hours (for slower growers the starter culture was used); and OD600 was measured, where OD600 = 1 was assumed to represent 8 × 10^9^ cells. 1 x 10^6^ cells of each non-producer strain were then plated onto Lennox media with each of the eight antibiotics at six concentrations (see Table S3).

Subsequently, all plates were incubated at 30 °C for 7 days and then photographed; our scoring was based on these images. We examined the set of 6 plates for each strain (10 producers and 10 non-producers) against each of the eight antibiotics (i.e. 6 x 8 plates per strain tested) for signs of growth (indicating resistance in the strain). We scored the minimal inhibitory concentration (MIC, the lowest concentration with no growth) and lowest effective concentration (LEC, the lowest concentration with an inhibitory effect) for each pairing of strain x antibiotic (see S4).

We then assigned each strain two scores: one based on the antibiotic concentration at which the MIC was reached, and one based on the LEC, on a scale from 1 (if the MIC or LEC was demonstrated at the lowest concentration of antibiotics above 0) to 5 (if the MIC or LEC occurred at the highest of the six concentrations used); we gave a score of 6 if the MIC or LEC was higher than all concentrations of antibiotic used.

**Table S3.** Antibiotic concentrations for MIC tests.

| Antibiotic | Concentrations Used | Target | Compound type |
| --- | --- | --- | --- |
| Chloramphenicol | 0, 2.5, 5, 10, 25, 50 µg/ml | Translation inhibitor | Synthetic |
| Rifampicin | 0, 0.5, 1, 2, 5, 10 µg/ml | Translation inhibitor | Polyketide |
| Streptomycin | 0, 5, 10, 20, 50, 100 µg/ml | Translation inhibitor | Aminoglycoside |
| Vancomycin | 0, 1, 2, 4, 10, 20 µg/ml | Cell wall synthesis inhibitor | Glycopeptide |
| Phosphomycin | 0, 5, 10, 20, 50, 100 µg/ml | Cell wall synthesis inhibitor | Small molecule |
| Nalidixic Acid | 0, 2.5, 5, 10, 25, 50 µg/ml | DNA gyrase inhibitor | Synthetic quinolone |
| Apramycin | 0, 5, 10, 20, 50, 100 µg/ml | Translation inhibitor | Aminoglycoside |
| Ampicillin | 0, 10, 20, 40, 100, 200 µg/ml | Cell wall synthesis inhibitor | β lactam |

## S4.

R markdown-format scripts, input datafiles, and html output files for the analyses in Figures 1, 2, and 3 are provided as a single R project folder and at https://github.com/dougwyu/screening_Innocent_et_al.

## S5. A human example of a successful screening mechanism

Any automobile-breakdown insurance company is faced with a hidden information problem. Potential clients differ in the probability that their cars will need rescue. Owners of poor-quality cars are more willing to purchase insurance but will impose higher costs on insurers with their more frequent callouts. Owners of high-quality cars are less willing to pay for insurance but would be more profitable to insurers. The challenge for insurers is to find a way to charge a higher price to owners of poor-quality cars and a lower price to owners of high-quality cars, without needing to inspect the cars.

One solution can be viewed at www.rac.co.uk/breakdown-cover (accessed 30 May 2018). Adding the ‘At home cover’ rescue option costs an additional £5/month, nearly doubling the cost of the cheapest cover at £5.50/month. Poor-quality cars run a nontrivial risk of failing to start in the morning. For high-quality cars, this risk is negligible. In screening terms, the absence of ‘At Home’ recovery is a burden that owners of high-quality cars are better able to endure. If priced and designed correctly, the two types of owners will voluntarily sign up for the two different coverage levels.

A similar design challenge applies to costly, honest signalling, in which the signal has to be costly in a way that reveals a specific hidden quality. For instance, an expensive car is good at signalling wealth and poor at signalling fidelity or niceness.

